# Aggregation of the constitutively active K296E rhodopsin mutant contributes to retinal degeneration

**DOI:** 10.1101/2025.03.26.643112

**Authors:** Sreelakshmi Vasudevan, Vivek Prakash, Paul S.–H. Park

## Abstract

A K296E mutation in rhodopsin causes autosomal dominant retinitis pigmentosa, a progressive retinal degenerative disease. Early *in vitro* characterizations of this mutation studied on a bovine rhodopsin background indicated that the mutation causes the receptor to be constitutively active. This molecular defect has been the primary focus when considering the pathogenic mechanism of the mutation. Knockin mice expressing the K296E rhodopsin mutant were generated and characterized to better understand the pathogenic mechanism of the mutation. Knockin mice exhibited progressive retinal degeneration characteristic of retinitis pigmentosa. The K296E rhodopsin mutant mislocalized in photoreceptor cells and, surprisingly, appeared to aggregate, as indicated by the dye PROTEOSTAT, which binds protein aggregates. The propensity of the K296E rhodopsin mutant to aggregate was tested and confirmed *in vitro* but was dependent on the species background of rhodopsin. The K296E mutation on either murine or human rhodopsin backgrounds exhibited similar propensities to aggregate. The same mutation on a bovine rhodopsin background, however, exhibited a lower propensity to aggregate, indicating this species background does not adequately model the effects of the K296E mutation. In contrast to previous expectations, we demonstrate here that aggregation of the K296E rhodopsin mutant can promote photoreceptor cell loss.

## INTRODUCTION

Rhodopsin is the light-activated G protein-coupled receptor expressed in rod photoreceptor cells of the retina. The rhodopsin gene is a hotspot for mutations, with over 100 mutations identified as a cause of retinitis pigmentosa (RP), a progressive retinal degenerative disease (1, 2). Rhodopsin mutations have been classified clinically according to the severity of the retinal degeneration phenotype and molecularly based on the type of defect promoted in the receptor (3–5). Mutations can cause a variety of molecular defects, with mutations causing receptor misfolding and aggregation forming the largest class of mutations (3). The P23H mutation was the first identified mutation in patients with autosomal dominant RP (adRP) (6), and it is the most extensively characterized mutation both *in vitro* and *in vivo*. This point mutation causes a moderate retinal degeneration phenotype and causes the receptor to misfold and aggregate (3, 4, 7).

Light activation of rhodopsin occurs via isomerization of 11-*cis* retinal, which is covalently linked to a lysine residue at position 296 of rhodopsin (Fig. 1A). Activation of rhodopsin is terminated through regulatory mechanisms involving the phosphorylation of the receptor by rhodopsin kinase and the binding of arrestin (8). A mutation of the lysine residue to glutamic acid (K296E) is a cause of adRP with a more severe retinal degeneration phenotype with earlier onset compared to the phenotype promoted by the P23H mutation (9, 10). Two types of molecular defects caused by the point mutation have been identified *in vitro*, however, only one has been demonstrated to occur *in vivo*. Initial *in vitro* characterization of the K296E rhodopsin mutant demonstrated that the mutation causes the receptor to be constitutively active (11). The constitutive activity of the K296E rhodopsin mutant was later demonstrated *in vivo* in a transgenic mouse model (12, 13). The constitutive activity of the K296E mutant, however, did not result in persistent activation of the phototransduction cascade because of regulatory mechanisms involving arrestin, but rather, resulted in stable interactions between the mutant and arrestin, which appears to contribute to the retinal degeneration phenotype (12–14). It has been unclear why some mutations causing constitutive activity in rhodopsin lead to retinal degeneration and are classified as a cause of RP whereas others lead to a more stationary form of night blindness and classified as a cause of congenital stationary night blindness (CSNB) (8).

**Figure 1.**
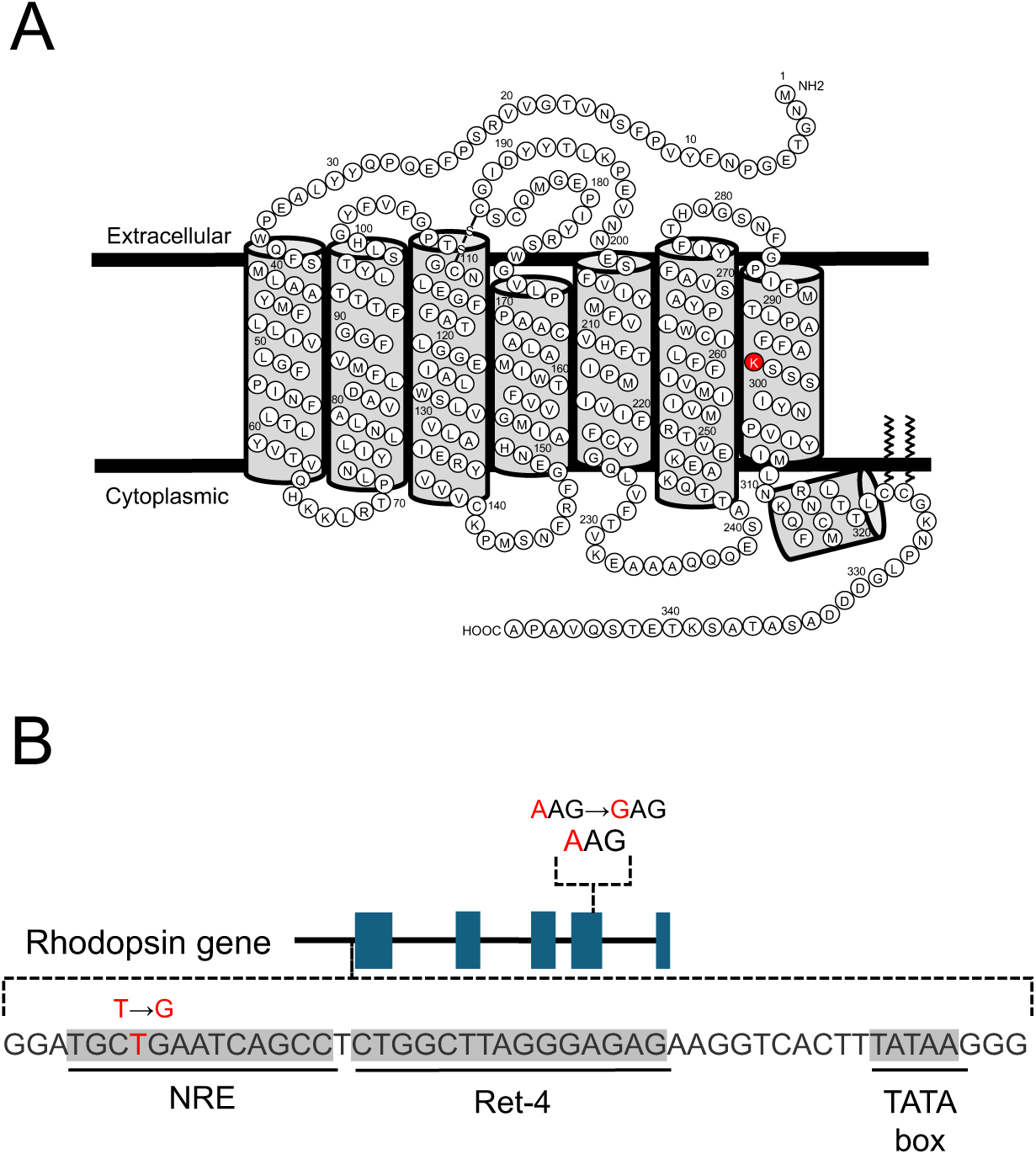
K296E mutation in rhodopsin. **A.** Secondary structure of murine rhodopsin is shown with the lysine residue at position 296 highlighted in red. **B.** The rhodopsin gene is illustrated highlighting the AAG (lysine) to GAG (glutamic acid) mutation in exon 4 present in K296E rhodopsin knockin mice and the promoter region shown highlighting a thymine (T) to guanine (G) base change within the NRE cis-regulatory element that is present in line K29-1.

Other *in vitro* characterizations have suggested the possibility that the K296E rhodopsin mutant can also misfold and aggregate (15, 16). This molecular defect, however, has not been demonstrated to occur *in vivo*. To determine whether the K296E rhodopsin mutation can promote misfolding and aggregation of the receptor and contribute to retinal degeneration, the mutant was characterized here both *in vitro* and *in vivo*. *In vitro* characterizations utilized a Förster resonance energy transfer (FRET)-based method in cells to detect aggregates of rhodopsin (17). For *in vivo* characterizations, a knockin mouse expressing the K296E rhodopsin mutant was generated by CRISPR/Cas9 gene editing method since previous *in vivo* studies utilized a transgenic mouse model (13). In the current study, we demonstrate that the K296E rhodopsin mutant aggregates both *in vitro* and *in vivo*, and that the aggregation of the receptor can contribute to the retinal degeneration phenotype.

## RESULTS

### Generation and initial characterization of K296E rhodopsin knockin mice

A knockin mouse that expresses the K296E rhodopsin mutant was generated by CRISPR/Cas9 gene editing methods to study the mutant *in vivo*. Knockin mice have proven to more accurately model adRP compared to transgenic mice, where the expression of mutants can be variable (18). A guide RNA was selected so that Cas9 endonuclease would cut the rhodopsin gene in exon 4 in the vicinity of the sequence corresponding to codon 296. Homology directed repair in the presence of a replacement oligonucleotide introduced the lysine (AAG) to glutamic acid (GAG) mutation at codon position 296 (Fig. 1B). Three founder mice (K29-1, K29-4, and K29-21) were identified that contained the AAG to GAG mutation and lines were established for each. The rhodopsin gene, including the promoter region, was sequenced for each of the three lines. No sequence changes were identified in K29-4 and K29-21 lines except for those introduced by the replacement oligonucleotide. Samples from the K29-1 line, however, exhibited a thymine (T) to guanine (G) base change within the NRE cis-regulatory element in the rhodopsin promoter region (Fig. 1B).

The retinal phenotype was characterized in each of these lines in both heterozygous (*Rho*^K296E/+^) and homozygous (*Rho*^K296E^) backgrounds. The loss of photoreceptor cells was quantified by counting the number of nuclei spanning the outer nuclear layer. All three lines exhibited the loss of photoreceptor cells, with the loss in *Rho*^K296E^ mice more severe than that in *Rho*^K296E/+^ mice (Fig. 2). The level of photoreceptor cell loss was less severe in the K29-1 line compared to the other two lines in both heterozygous and homozygous backgrounds. The K29-4 and K29-21 lines exhibited similar levels of photoreceptor cell loss. Thus, the mutation in the promoter region present in the K29-1 line appears to diminish the effect of the mutation.

**Figure 2.**
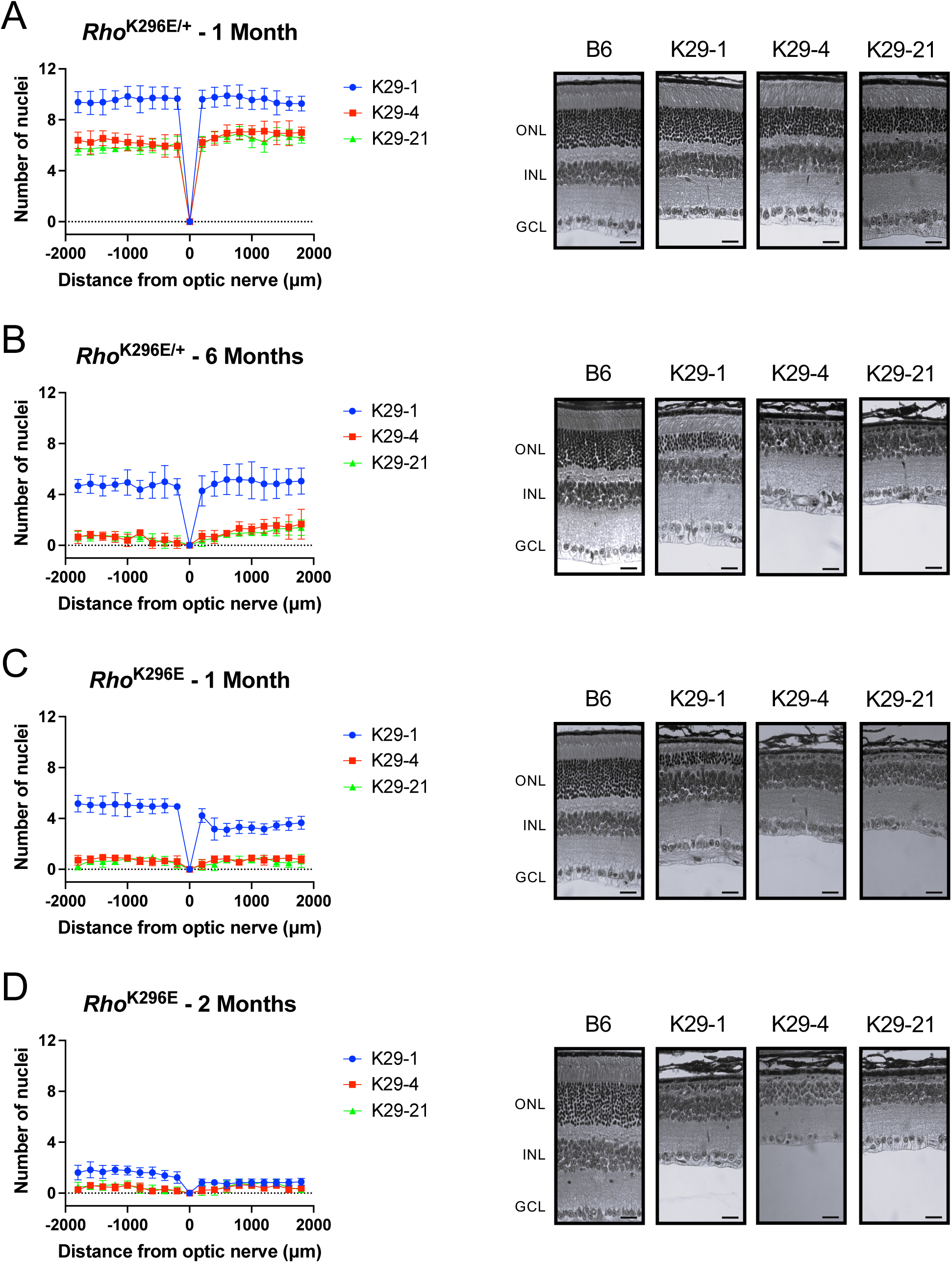
Photoreceptor cell loss in K296E rhodopsin knockin mice. Spider plots quantifying the number of photoreceptor cell nuclei in the inferior (negative) and superior (positive) regions of the retina of 1-month-(A) and 6-month-old (B) *Rho*^K296E/+^ mice or 1-month-(C) and 2-month-old (D) *Rho*^K296E^ mice are shown on the left-hand side. Data are shown for K29-1, K29-4, and K29-21 lines and age of mice are indicated. The mean and standard deviation are shown at different distances from the optic nerve (*n* = 6). Corresponding images of retinal sections are shown on the right-hand side for B6 mice and the K29-1, K294, and K29-21 lines. The outer nuclear layer (ONL), inner nuclear layer (INL), and ganglion cell layer (GCL) are labeled. Scale bar, 25 μm.

### Expression of rhodopsin in K296E rhodopsin knockin mice

To determine if the mutation in the promoter region present in the K29-1 line impacts the expression of rhodopsin, RT-qPCR and Western blot analysis was conducted on retinal samples from 2-week-old heterozygous and homozygous mice to quantify the levels of rhodopsin transcript and protein. Rhodopsin transcripts were normalized to transcripts of *18s rRNA*, which does not consider any photoreceptor cell loss, and *Gnat1*, which does take into account photoreceptor cell loss. In heterozygous mice, the level of rhodopsin transcripts was similar to B6 mice regardless of normalization to *18s rRNA* or *Gnat1* for all three mouse lines (Fig. 3A), which indicates that loss of photoreceptor cells is minimal at this age. The level of rhodopsin protein, as assessed by Western blot, was a little less than half of that in B6 mice and was similar for all three lines (Fig. 3B). Thus, most of the K296E rhodopsin mutant may be degraded in photoreceptor cells like the P23H and G188R rhodopsin mutants (7).

**Figure 3.**
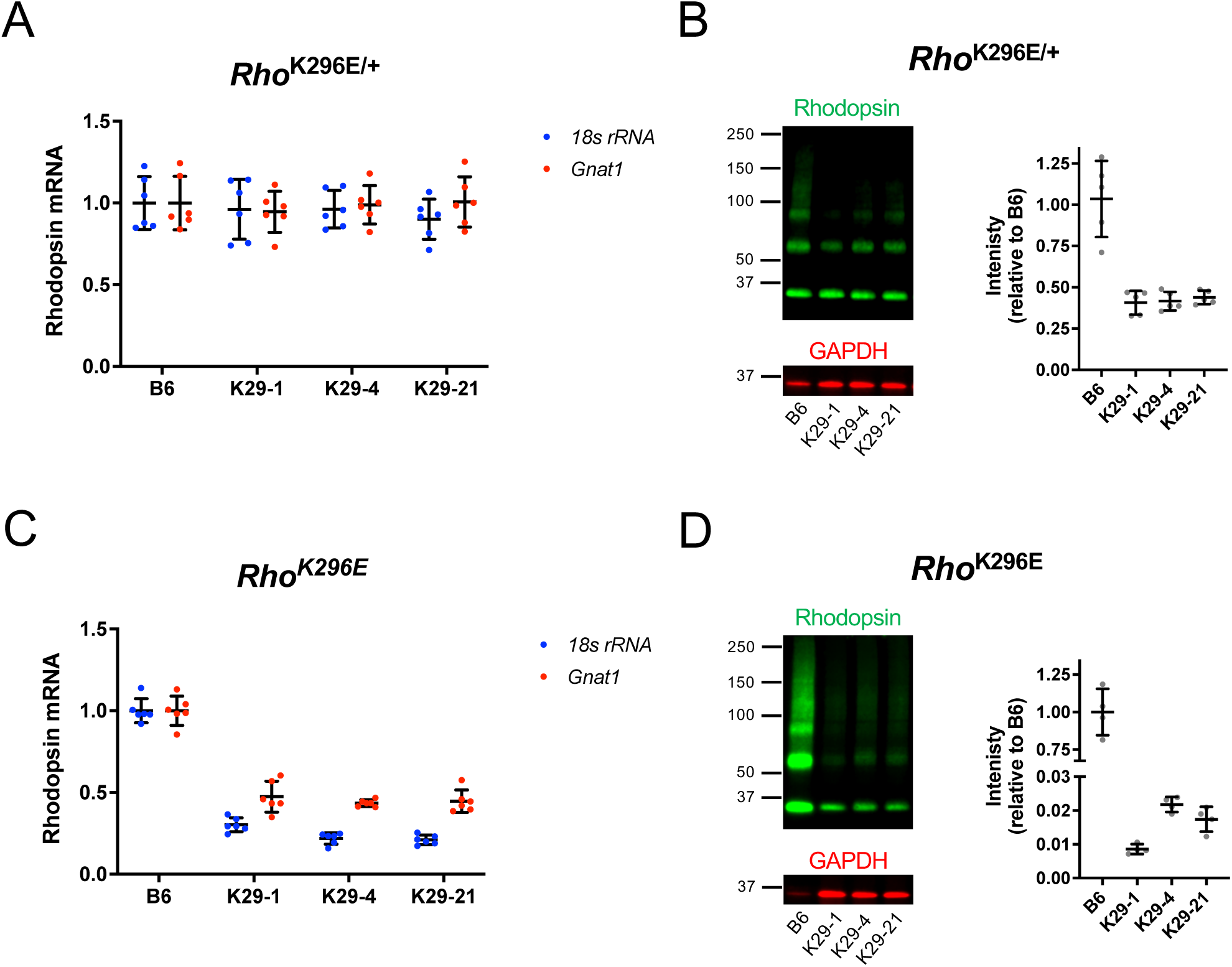
Rhodopsin expression in K296E rhodopsin knockin mice. Rhodopsin transcripts and protein were quantified from the retina of 2-week-old B6 and *Rho*^K296E/+^ and *Rho*^K296E^ mice from lines K29-1, K29-4, and K29-21. **A, C.** Rhodopsin transcripts in *Rho*^K296E/+^ (A) and *Rho*^K296E^ (C) mice were quantified by RT-qPCR and expressed relative to the mean value for B6 mice and normalized to *18s rRNA* (blue) or *Gnat1* (red). Individual data points are shown and are plotted along with the mean and standard deviation (*n* = 6). **B, D.** Rhodopsin protein in *Rho*^K296E/+^ (B) and *Rho*^K296E^ (D) mice was quantified from Western blots of retinal extracts labeled with anti-1D4 (green) or anti-GAPDH (red) antibodies. Molecular weight markers are indicated in kDa. The intensity of bands in Western blots corresponding to rhodopsin were normalized to the intensity of the band corresponding to GAPDH and expressed relative to the mean value for B6 mice. Individual data points are shown along with the mean and standard deviation (*n* = 6). Tukey’s post-hoc analysis indicated that the level of rhodopsin in *Rho*^K296E^ mice of line K29-1 was significantly different than that in lines K29-4 and K29-21 (P < 0.05).

In homozygous mice, rhodopsin transcript levels were lower compared that in B6 mice and all three lines exhibited similar levels of rhodopsin transcripts (Fig. 3C). Homozygous mice did exhibit differences in rhodopsin transcript levels when normalized to *18s rRNA* or *Gnat1*, indicating that there is a loss of photoreceptor cells at this age. Even when photoreceptor loss was taken into account by normalization to *Gnat1*, rhodopsin transcript levels were a little under half of that for B6 mice, indicating that transcription is affected in these mutant mice. While 2-week-old homozygous mice from line K29-4 exhibits significant photoreceptor cell loss, photoreceptor loss is not apparent in homozygous mice from line K29-1 (Supplementary Fig. 1 and Fig. 4B). Thus, retinal degeneration may affect transcription of rhodopsin in line K29-4, as was the case in homozygous mice expressing the P23H or G188R rhodopsin mutants (7). The lower level of rhodopsin transcripts in line K29-1 cannot be explained by retinal degeneration but may be a result of the mutation in the rhodopsin promoter region. The level of rhodopsin mutant protein expressed in homozygous mice was about 2 % compared to that in B6 mice for lines K29-4 and K29-21 and only 1 % for line K29-1 (Fig. 3D). Thus, most of the mutant appears to be degraded, and line K29-1 appears to express the mutant at half the level of that in lines K29-4 and K29-21. Taken together, the mutation in the promoter region of line K29-1 appears to reduce the expression of the K296E rhodopsin mutant.

**Figure 4.**
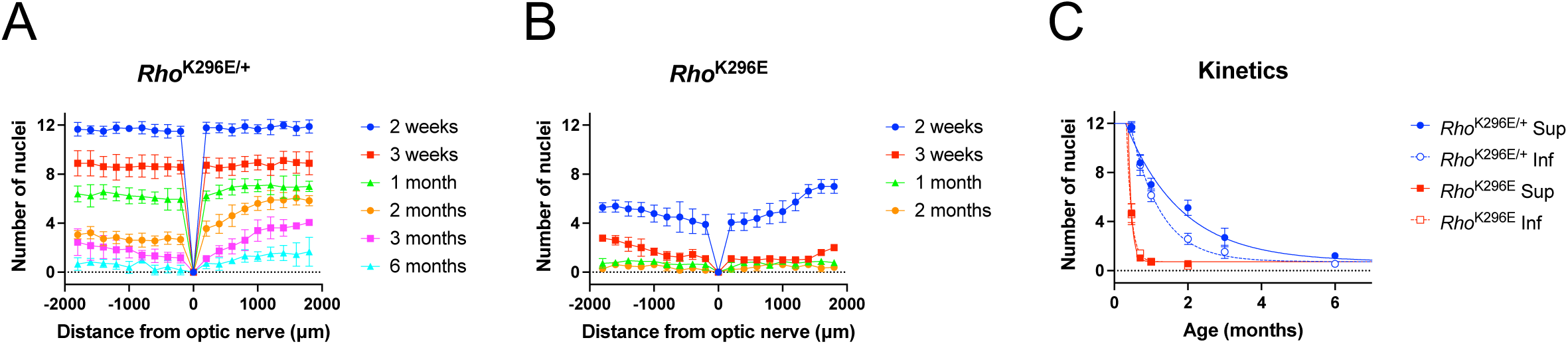
Progressive photoreceptor cell loss in *Rho*^K296E/+^ and *Rho*^K296E^ mice. A, **B.** Spider plots. The number of photoreceptor cell nuclei in the inferior (negative) and superior (positive) regions of the retina were quantified in retinal section images (e.g, Fig. 2) from *Rho*^K296E/+^ (A) and *Rho*^K296E^ (B) mice at the indicated ages. The mean and standard deviation are shown at different distances from the optic nerve (*n* = 6). **C.** Kinetics of photoreceptor cell loss. Mean values of the number of photoreceptor cell nuclei, and their associated standard deviation, are plotted as a function of age in the superior or inferior region of the retina in *Rho*^K296E/+^ (blue) and *Rho*^K296E^ (red) mice (*n* = 6). Data were fit by non-linear regression to determine the rate constants, which are reported in Supplementary Table 1.

### Characterization of photoreceptor cell loss promoted by K296E mutant rhodopsin

The K29-4 line was investigated from this point on to better understand the effect of the K296E rhodopsin mutation in photoreceptor cell loss. Spider plots of the number of nuclei spanning the outer nuclear layer were generated for 2 week – 6-month-old *Rho*^K296E/+^ mice (Fig. 4A) and 2 week – 2-month-old *Rho*^K296E^ mice (Fig. 4B). The degeneration was more severe in the inferior retina in *Rho*^K296E/+^ mice compared to that in the superior retina. The kinetics of the photoreceptor cell loss was determined for the central region of the superior and inferior retina (Fig. 4C). The rate of photoreceptor cell loss was about 2-fold faster in the inferior retina of *Rho*^K296E/+^ mice compared that in the superior retina. In contrast, the rate of photoreceptor cell loss was similar in the superior and inferior retina of *Rho*^K296E^ mice. The rate of photoreceptor cell loss in *Rho*^K296E^ mice was 6-fold faster or more compared to that in *Rho*^K296E/+^ mice.

### Mislocalization and aggregation of K296E mutant rhodopsin in photoreceptor cells

The localization of rhodopsin within photoreceptor cells was characterized by immunohistochemistry in 2-week-old mice using the anti-4D2 antibody (Fig. 5A), which detects the amino terminal region of rhodopsin (19). B6 mice exhibited staining by the anti-4D2 antibody only in the rod outer segment, demonstrating that WT rhodopsin is properly targeted without mislocalization. In contrast, the anti-4D2 antibody detected rhodopsin in both outer segments and mislocalized in the outer nuclear in both *Rho*^K296E/+^ and *Rho*^K296E^ mice. The outer segments were severely shortened in mutant mice, especially in *Rho*^K296E^ mice. Since only the mutant is expressed in *Rho*^K296E^ mice, a small fraction of K296E mutant rhodopsin appears to traffic to the rod outer segment, which was not previously observed for misfolding mutants of rhodopsin (7).

**Figure 5.**
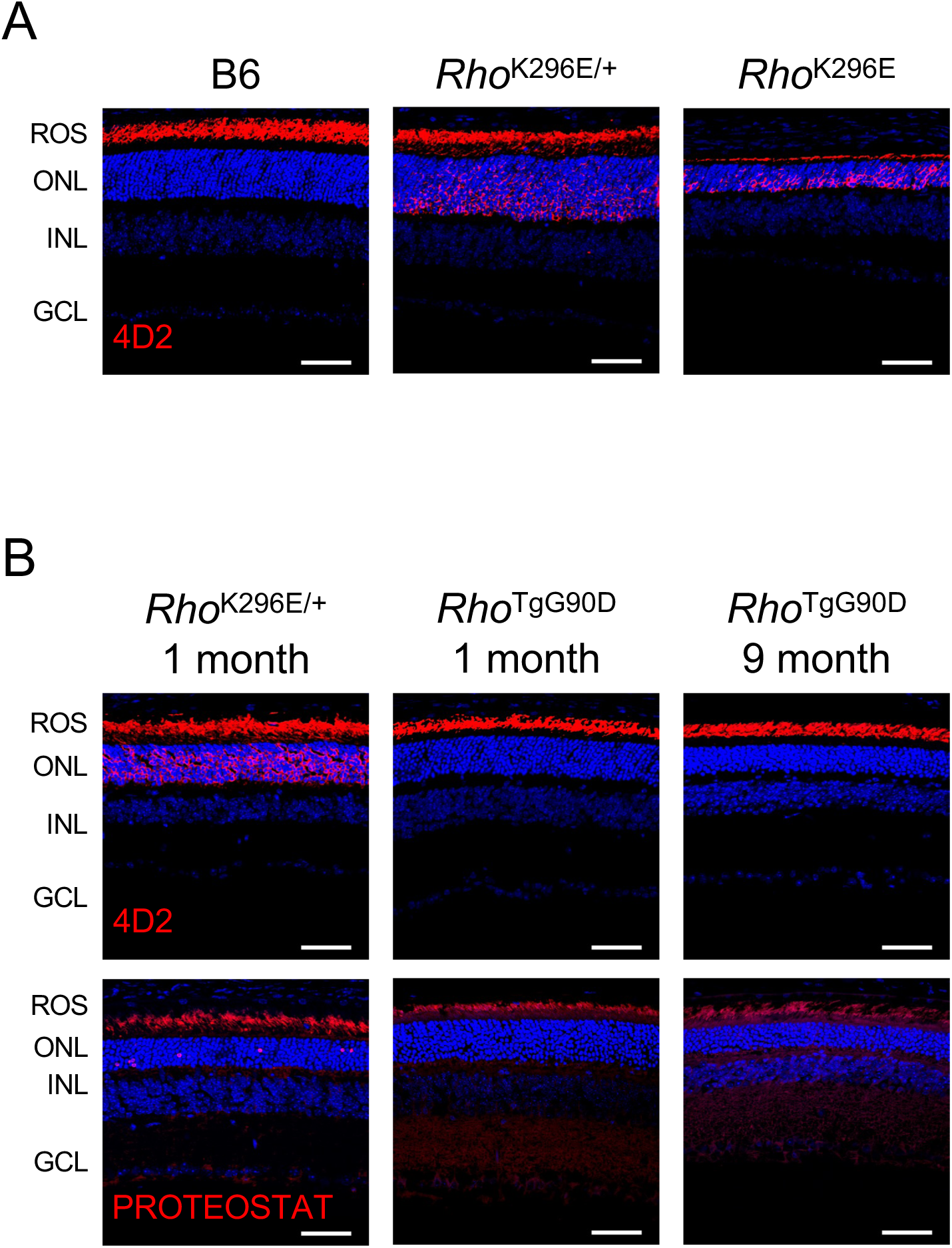
Mislocalization and aggregation of the K296E rhodopsin mutant. **A.** Confocal microscopy image of retinal cryosections from 2-week-old B6, *Rho*^K296E/+^ and *Rho*^K296E^ mice labeled with the anti-4D2 antibody (red). **B.** Confocal microscopy image of retinal cryosections from 1-month-old *Rho*^K296E/+^ mice or 1-month-and 9-month-old *Rho*^TgG90D^ mice labeled with the anti-4D2 antibody or PROTEOSTAT (red). Nuclei are labeled with DAPI (blue). Confocal microscopy images of retinal cryosections were obtained at 40× magnification. Rod outer segments (ROS), outer nuclear layer (ONL), inner nuclear layer (INL), and ganglion cell layer (GCL) are labeled. Scale bar, 50 μm.

Mislocalization of misfolding mutants of rhodopsin is accompanied by aggregation of the mutants, which can be detected in retinal cryosections by the dye PROTEOSTAT (7, 20), a molecular rotor dye that detects aggregated proteins (21). To determine if the mislocalization of the K296E rhodopsin mutant is accompanied by aggregation of the mutant, retinal cryosections from *Rho*^K296E/+^ mice were labeled with PROTEOSTAT. PROTEOSTAT labeling was observed in the outer nuclear layer, indicating that the mutant aggregates (Fig. 5B). Higher magnification images of the outer nuclear layer revealed that PROTEOSTAT labeling surrounded photoreceptor cell nuclei (Fig. 6A), which was the same labeling pattern displayed for misfolding mutants of rhodopsin that aggregate (7, 20). PROTEOSTAT-positive nuclei were both relatively healthy with a single large central chromocenter and unhealthy with disrupted nuclei. Thus, the K296E rhodopsin mutant mislocalizes in photoreceptor cells and aggregates similarly as previously characterized misfolding mutants of rhodopsin. The appearance of PROTEOSTAT labeling surrounding photoreceptor cell nuclei can precede the deterioration of the nuclei.

**Figure 6.**
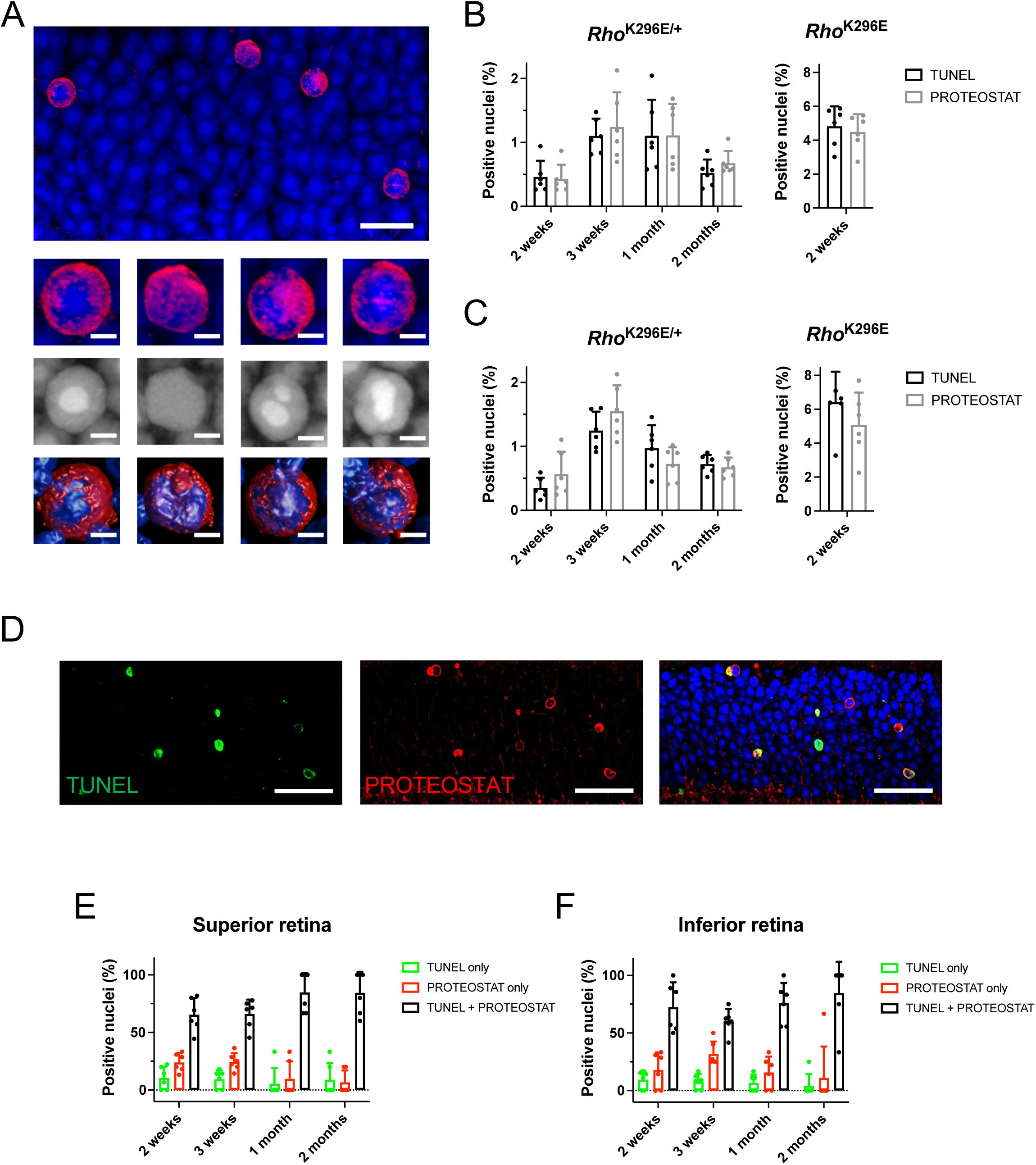
Relationship between PROTEOSTAT-and TUNEL-positive photoreceptor cells. **A.** Maximum intensity projection image of retinal cryosection from 3-week-old *Rho*^K296E/+^ mice labeled with PROTEOSTAT (red) and NucBlue (blue). Scale bar, 5 μm. Maximum intensity projection image was generated using images obtained at 100× magnification with 2.5× digital zoom with deconvolution. Zoomed in images of PROTEOSTAT-positive nuclei are shown below including grey scale image of NucBlue staining only and surface rendered image. Scale bar, 1 μm. **B, C.** The number of nuclei labeled by TUNEL (black) or PROTEOSTAT (gray) in the outer nuclear layer were quantified in the superior (B) and inferior (C) regions of the retina in *Rho*^K296E/+^ and *Rho*^K296E^ mice at the ages indicated. Individual data points are plotted along with the mean and standard deviation (*n* = 6). **D.** Confocal microscopy image of retinal cryosections from 3-week-old *Rho*^K296E/+^ mice labeled with both PROTEOSTAT (red) and TUNEL (green). Nuclei are labeled with NucBlue (blue). Confocal microscopy images were obtained at 100× magnification and both separated and merged images are shown. Scale bar, 25 μm. **E, F.** The number of photoreceptor cell nuclei labeled by TUNEL (green) or PROTEOSTAT (red) alone or co-labeled with TUNEL and PROTEOSTAT (black) were quantified from confocal microscopy images of retinal cryosections taken from *Rho*^K296E/+^ mice at ages indicated (e.g., Fig. 6D). Individual data points are plotted along with the mean and standard deviation (*n* = 6).

To determine if aggregation is a common feature of other constitutively active rhodopsin mutants, a transgenic mouse expressing the G90D rhodopsin mutant on a null rhodopsin background (*Rho*^TgG90D^) was examined (22). The G90D rhodopsin mutant is constitutively active and is classified as a cause of CSNB but can also cause mild retinal degeneration (23–26).

Labeling retinal cryosections with the anti-4D2 antibody did not detect any mislocalization of rhodopsin. Likewise, no PROTEOSTAT-positive photoreceptor cell nuclei were detected. Thus, not all constitutively active rhodopsin mutants cause aggregation and photoreceptor cell death caused by constitutively active rhodopsin mutants can have different molecular origins.

### Relationship between aggregation and photoreceptor cell death

To determine if the PROTEOSTAT-labeled aggregates contribute to photoreceptor cell death, the relationship between aggregation, detected by PROTEOSTAT, and photoreceptor cell death, assessed by TUNEL, was determined. The level of TUNEL and PROTEOSTAT positive photoreceptor cell nuclei were quantified in the retina of mice at different ages. In *Rho*^K296E/+^ mice, the level of TUNEL and PROTEOSTAT positive photoreceptor cell nuclei appeared to be correlated with the peak occurring at about 3 weeks of age (Figs. 6B and 6C). In *Rho*^K296E^ mice, where photoreceptor cell loss is more severe, the level of TUNEL and PROTEOSTAT positive nuclei was higher than that observed in *Rho*^K296E/+^ mice. Thus, the level of photoreceptor cell death is related to the appearance of PROTEOSTAT-labeled photoreceptor cell nuclei. To determine further the relationship between photoreceptor cell death and aggregation, the level of colabeling of photoreceptor cell nuclei by TUNEL and PROTEOSTAT was determined (Fig. 6D). At all ages of *Rho*^K296E/+^ mice tested, a majority of nuclei were colabeled by TUNEL and PROTEOSTAT whereas only a relatively few nuclei were labeled by TUNEL only (Figs. 6E and 6F). Up to a quarter of nuclei were labeled only by PROTEOSTAT. Taken together, it appears that the appearance of aggregates surrounding photoreceptor cell nuclei can precede cell death and that these aggregates can contribute to the onset of cell death.

### In vitro characterization of rhodopsin K296E mutant aggregates

To examine the propensity of K296E rhodopsin to aggregate and determine whether this is a species-specific property, a FRET-based method we developed to detect rhodopsin aggregates in HEK293 cells was used (17). The K296E mutation was introduced on the murine, human, and bovine rhodopsin backgrounds. WT and mutant rhodopsins were tagged with either a yellow fluorescent protein (YFP) variant or mTurquoise2 (mTq2) (27, 28) and coexpressed in HEK293 cells. WT rhodopsin typically forms oligomers in photoreceptor cells (29). The formation of oligomers and aggregates can be differentiated in our assay by the sensitivity of the FRET signal to the mild detergent *n*-dodecyl-*β*-D-maltoside (DM), where DM-sensitive and DM-insensitive FRET derive from oligomers and aggregates, respectively. DM-sensitive and DM-insensitive FRET was measured for WT and mutant rhodopsins (Figs. 7A and 7B). Only the specific FRET signal was considered since the physiological relevance of the FRET signal equal to or less than non-specific FRET is ambiguous (17).

**Figure 7.**
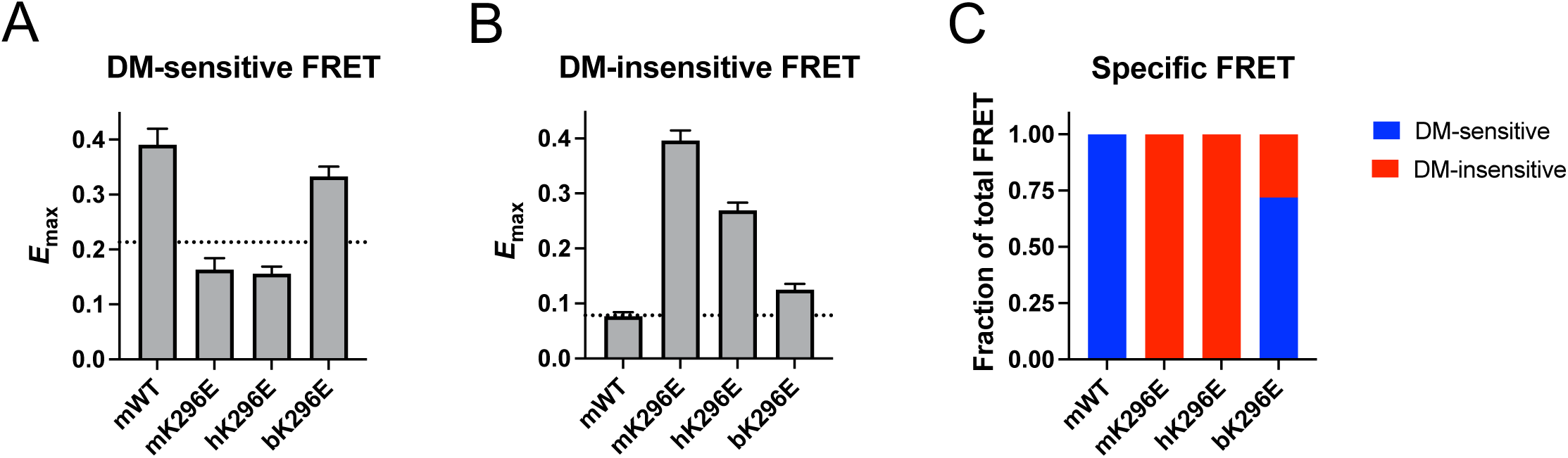
Characterization of A, **B.** FRET was conducted on HEK239 cells expressing mTq2-and YFP-tagged WT murine rhodopsin or murine (mK296E), human (hK296E), or bovine (bK296E) rhodopsin with the K296E mutation. Fitted values of the maximal FRET efficiency (*E*_max_) and the standard errors from the fits are shown for DM-sensitive (A) and DM-insensitive (B) components of generated FRET curves (Supplementary Fig. 2). The non-specific *E*_max_, defined previously (44), is indicated by the dotted lines. All fitted parameters are reported in Supplementary Table 2. **C.** The fraction of the total FRET signal derived from DM-sensitive (blue) and DM-insensitive (red) FRET is shown for each of the rhodopsins examined. Only the specific FRET signal was considered.

For murine WT rhodopsin, only specific DM-sensitive FRET was detected, indicating that rhodopsin predominantly forms oligomers rather than aggregates in HEK293 cells (Fig. 7C), as shown previously (7, 30). For the K296E mutation on both murine and human rhodopsin backgrounds, only specific DM-insensitive FRET was detected, indicating that the mutants predominantly form aggregates in HEK293 cells. In contrast, the K296E mutation on a bovine rhodopsin background predominantly displayed specific DM-sensitive FRET with a small DM-insensitive FRET signal, indicating the mutant forms mostly oligomers and some aggregates.

These *in vitro* studies demonstrate the propensity of the K296E mutation to promote rhodopsin aggregation is dependent on the species. Both murine and human K296E rhodopsin mutants are predicted to aggregate, which is consistent with studies in the K296E mutant rhodopsin knockin mice.

## DISCUSSION

Characterization of knockin mice expressing the K296E rhodopsin mutant has clarified two major questions about the K296E mutation. The first question is what is the molecular cause of photoreceptor cell death promoted by the mutation? The second question is why some constitutively active mutations cause RP whereas others cause CSNB? The predominant focus of the pathogenic mechanism of the K296E mutation has been on the effect or consequence of the constitutive activity of the receptor promoted by the mutation (12, 13). We demonstrate here both *in vivo* and *in vitro* that the K296E rhodopsin mutant aggregates, and that aggregation can contribute to photoreceptor cell death.

Many early *in vitro* characterizations of the K296E mutation were conducted on a bovine rhodopsin background. The earliest study demonstrated that the mutant is constitutively active (11), which is a reason why this defect has been the primary focus when considering the pathogenic mechanism of the mutation. Later *in vitro* studies showed that while a large fraction of the K296E rhodopsin mutant can fold properly, there were also indications that it may also misfold and aggregate (16, 31). We demonstrate here that the bovine rhodopsin background is a poor molecular model to examine the effect of mutation in rhodopsin to understand human disease. The K296E rhodopsin mutation promoted similar aggregation profiles on the murine and human rhodopsin backgrounds, where the mutants were predominantly aggregated (Fig. 7C).

The same mutation on the bovine rhodopsin background, however, resulted in a less severe aggregation profile where most of the mutant formed oligomers like it does in WT rhodopsin. We demonstrated previously that the aggregation profile of the P23H mutation on a bovine rhodopsin background is also less severe than on the human or murine rhodopsin backgrounds (7, 32). While the molecular interactions stabilizing the structures of WT murine and bovine rhodopsin appear to be conserved (33), the impact of mutations differ on the two backgrounds. So far, it appears that *in vitro* properties of mutations in murine and human rhodopsin backgrounds are equivalent, thereby justifying the use of knockin mouse models to study rhodopsin-mediated adRP.

The K296E rhodopsin mutation shares some common properties as the P23H and G188R rhodopsin mutations, which are classified as mutations that cause receptor misfolding and aggregation leading to adRP (3, 7). *Rho*^K296E/+^ mice like the *Rho*^P23H/+^ and *Rho*^G188R/+^ mice display progressive retinal degeneration where the inferior retina exhibits a more rapidly progressing degeneration compared to the superior retina (Figs. 4A and 4C). In knockin mice, all three mutants are shown to mislocalize in the photoreceptor cell and aggregate (Fig. 5B), as demonstrated by PROTEOSTAT labeling that surrounds the nuclei (Fig. 6A). This labeling of photoreceptor cell nuclei has been demonstrated to derive from the aggregation of mutant rhodopsin (7). Aggregation seems to be a major driver of photoreceptor cell death in mice expressing the K296E rhodopsin mutant since there were relatively few photoreceptor cells that were only TUNEL positive while most exhibited PROTEOSTAT labeling either alone or together with TUNEL labeling (Figs. 6D-6F). This pattern is also observed in in *Rho*^P23H/+^ and *Rho*^G188R/+^ mice where aggregation is a major driver of photoreceptor cell death (20).

There are key differences exhibited by the K296E rhodopsin mutant that suggest aggregation may not be the sole source of photoreceptor cell loss. The level of both TUNEL and PROTEOSTAT positive cells is lower in mice expressing the K296E rhodopsin mutant compared to those expressing either the P23H or G188R rhodopsin mutants (7). Despite the lower levels of TUNEL and PROTEOSTAT positive cells, the rate of photoreceptor cell loss in *Rho*^K296E/+^ mice is similar to *Rho*^G188R/+^ mice and faster than that in the *Rho*^P23H/+^ mice (Supplementary Fig. 3). A variety of cell death mechanisms have been proposed for rhodopsin-mediated photoreceptor cell death (34). TUNEL staining is indicative of apoptosis, but other cell death mechanisms may also be detected by TUNEL albeit with variable efficiency (35). Since TUNEL positive cells are lower in *Rho*^K296E/+^ mice but the rate of photoreceptor cell loss is similar or greater than in *Rho*^P23H/+^ and *Rho*^G188R/+^ mice, this discrepancy may point to an additional cell death mechanism than that promoted by rhodopsin aggregation with different kinetics or TUNEL staining efficiency.

The additional cell death mechanism may be a byproduct of the originally attributed constitutive activity promoted by the K296E rhodopsin mutation, which results in stable interactions with arrestin (12, 13). The constitutively active mutant would fold properly and be capable of being transported to the rod outer segment with some being mislocalized due to stable interactions formed with arrestin (12). In *Rho*^K296E^ mice, there is evidence that some of the mutant can traffic to the rod outer segment (Fig. 5A). Moreover, in Western blots of retinal extracts from *Rho*^K296E^ mice, there is a band corresponding to the monomeric receptor species (Fig. 3D). Previously, we demonstrated that G188R and P23H rhodopsin mutants migrate as dimeric or larger species in Western blots indicative of aggregation. Thus, there appears to be a small population of K296E rhodopsin mutant that is properly folded and likely constitutively active.

Studies here also provide an explanation on why different constitutively active mutants are classified as a cause of different diseases. One class of constitutively active mutations (G90V, S186W, D190N, K296, K296M) are classified as a cause of RP, which causes a more severe retinal degeneration phenotype, whereas another class of constitutively active mutations (G90D, T94I, A292E, A295V) are classified as a cause of CSNB, which causes a milder phenotype of night blindness originally thought to be without retinal degeneration (8). It has been shown later that the G90D rhodopsin mutation can indeed cause mild retinal degeneration in both mice and patients (25, 26). Nonetheless, we demonstrate here that the G90D rhodopsin mutant causes retinal degeneration in an aggregation independent manner (Fig. 5B). Thus, the distinction between constitutively active mutants classified as a cause of RP versus those that cause CSNB may be the ability of the mutants to form aggregates for the former. In support of this idea, the constitutively active D190N rhodopsin mutant has also previously been shown to form aggregates detectable by PROTEOSTAT (36).

The severity of the retinal degeneration phenotype appears to be related to the expression level of the K296E rhodopsin mutant. The thymine to guanine nucleotide mutation in the NRE *cis*-regulatory element of the rhodopsin promoter in line K29-1 (Fig. 1B) is predicted to decrease the expression of rhodopsin transcripts based on an *in vivo* reporter-based assay using a minimal rhodopsin promoter (37). While the expression of the K296E rhodopsin mutant does appear to be affected by the mutation in the promoter region, the magnitude of the effect is unclear. In homozygous mice of line K29-1, expression of the mutant is affected (Fig. 3D), although effects at the level of transcription are obscured because of retinal degeneration in the other lines (Fig. 3C). In heterozygous mice of line K29-1, no difference is observed in the expression of rhodopsin at both transcript and protein levels (Figs. 3A and 3B), which may indicate the presence of compensatory mechanisms and a more modest change in the expression of rhodopsin due to mutation in the promoter region. Regardless of the magnitude of the effect of the mutation in the promoter region on the expression of the K296E rhodopsin mutant, most of the mutant is degraded as it is in knockin mice expressing the P23H and G188R rhodopsin mutants (7). The amount of mutant present in photoreceptor cells is minimal compared to WT rhodopsin, yet this small population can aggregate and cause photoreceptor cell death. Thus, relatively small changes in the expression of the mutant can change the severity of the phenotype.

In summary, we demonstrate here that the constitutively active K296E rhodopsin mutation causes aggregation of the receptor, which appears to promote photoreceptor cell death. Over half of rhodopsin mutations causing adRP have already been shown to cause receptor misfolding and aggregation and it now appears there are even more since mutations classified in other categories may also cause misfolding and aggregation. We demonstrate that relatively small changes in expression of the mutant may have comparatively large effects on the retinal degeneration phenotype, which indicates even small therapeutic changes can have profound effects in prolonging vision. Lastly, studies here indicate that the differentiation of constitutively active rhodopsin mutants as a cause of RP or CSNB may be based on whether the mutation can also promote receptor aggregation.

## MATERIALS AND METHODS

### Mice

All animal studies reported here were conducted using protocols approved by the Institutional Animal Care and Use Committee at Case Western Reserve University School of Medicine. Both male and female mice were used for experiments. *Rho*^K296E^ mice were generated using CRISPR/Cas9 gene targeting at the Case Transgenic and Targeting Facility of Case Western Reserve University School of Medicine (Cleveland, OH). Fertilized embryos from C57Bl/6J mice were injected with Cas9 nuclease (PNA Bio, Thousand Oaks, CA), sgRNA with the sequence 5’GAGCTCTTAGCAAAGAAAGC (PNA Bio, Thousand Oaks, CA) and ssDNA replacement oligonucleotide with the sequence 5’CACCCACCAGGGCTCCAACTTCGGCCCCATCTTCATGACTCTGCCA**<u>GCA</u>**TTCTTTG CT**<u>GAG</u>**AGCTCTTCCATCTATAACCCGGTCATCTACATCATGTTGAACAAGCAGGTGCC TGGGCT (Integrated DNA Technologies, Coralville, IA), which contained the lysine (AAG) to glutamic acid (GAG) mutation and introduces a conservative substitution for alanine (GCT to GCA). Deep sequencing was conducted on the MiSeq System (Illumina, San Diego, CA) by the Genomics Core at Case Western Reserve University School of Medicine (Cleveland, OH) on samples from mosaic founder mice to identify mice with the desired mutation. 3 founder mice harboring the mutation were identified and each were backcrossed with C57Bl/6J mice for 10 generations to establish each line. A 10,000 base pair region of the genome containing the rhodopsin gene and promoter region in these mice was sequenced by PCR-amplifying overlapping fragments to confirm that mice only exhibited changes introduced in the replacement oligonucleotide. C57Bl/6J mice were purchased from The Jackson Laboratory (Bar Harbor, ME). Transgenic mice that were homozygous for the mutant G90D rhodopsin transgene on a null rhodopsin background (*Rho*^TgG90D^) were kindly provided by Dr. Paul Sieving (UC Davis, Sacramento, CA) (22).

### Quantifying photoreceptor cell loss

Hematoxylin and eosin (H&E)-stained retinal sections were prepared by Excalibur Pathology (Norman, OK), imaged on an Axio Scan.Z1 Slide Scanner equipped with a Hitachi HV-F203 camera and a Plan Apo 20×/0.8-NA objective (Carl Zeiss Microscopy, White Plains, NY) or a Leica DME compound microscope equipped with an EC3 digital camera and 40×/0.65-NA objective (Leica Microsystems, Buffalo, NY), and the number of nuclei spanning the outer nuclear layer quantified and analyzed as described previously (7, 38). Kinetics of photoreceptor cell loss were determined by fitting data by non-linear regression to a plateau followed by one phase decay equation in Prism 10 (GraphPad Software, San Diego, CA): *y = if(x < x_0_, y_0_, plateau + (y_0_ − plateau) × e^-k(-x^_0_^+x^*). The variable *y*_0_ was fixed at 12 and *plateau* was set to be common among all data. Fitted values of the rate constant (*k*) are reported with standard error of the fit.

### Rhodopsin expression

RT-qPCR was conducted on the LightCycler 96 Real-Time PCR System (Roche Diagnostics, Indianapolis, IN) to quantify rhodopsin transcripts in retinal extract. Sample preparation, primers for rhodopsin, transducin, and 18s rRNA transcripts, and qPCR procedures and analyses are the same those described previously (7, 25). Rhodopsin protein levels in retinal extracts of mice were quantified by Western blot analysis. Preparation of retina samples, SDS-PAGE using Novex 4-12% Tris-glycine gels (Invitrogen, Camarillo, CA), Western blotting procedures, and quantification of bands on Western blots by the Odyssey Fc Imaging System (LI-COR Biosciences, Lincoln, NE) were performed, as described previously (7). Primary antibodies against rhodopsin (anti-1D4) (39) and GAPDH (Cat. No. 10494-1-AP; Proteintech, Rosemont, IL) and IRDye 800CW donkey anti-mouse (Cat. No. 926-32212) or IRDye 680LT donkey anti-rabbit (Cat. No. 925-68023) secondary antibodies (LI-COR Biosciences, Lincoln, NE) were used.

### Labeling of retinal cryosections and confocal microscopy

Retinal cryosection preparation, immunohistochemistry, TUNEL assay, PROTEOSTAT labeling, confocal microscopy, and analysis and processing of images were conducted essentially as described previously (7, 20, 40). Rhodopsin was labeled with an anti-4D2 (Cat. No. MABN15, MilliporeSigma, Burlington, MA) primary antibody and Alexa Fluor 647 goat anti-mouse secondary antibody (Cat. No. A21237, Thermo Fisher Scientific, Waltham, MA). TUNEL assay was conducted using the One-step TUNEL In Situ Apoptosis Kit (Cat. No. E-CK-A324, Elabscience, Houston, Tx). PROTEOSTAT labeling was conducted using the PROTEOSTAT Aggresome Detection Kit (Enzo Life Sciences, Farmingdale, NY).

Confocal microscopy was performed on an Olympus FV1200 IX83 laser scanning confocal microscope (Evident Scientific, Waltham, MA) using either a UPlanXApo 40×/1.40 NA oil objective or UPLXAPO 100×/1.45 NA objective. Labeled cryosections were cover-slipped with DAPI Fluoromount-G mounting media (Southern Biotech, Birmingham, AL) for 40× imaging or with ProLong Glass Antifade Mountant with NucBlue stain (Invitrogen, Carlsbad, CA) for 100× imaging. DAPI and NucBlue were detected with 405 nm diode laser excitation and 425-460 nm emission, Alexa Fluor 647 and TUNEL positive cells were detected with 635 nm diode laser excitation and 655-755 nm emission, and PROTEOSTAT dye was detected by 559 nm diode laser excitation and 575-620 nm emission. Deconvolution, maximum projection image generation, and surface rendering were performed in Huygens Essential 23.10 software (Scientific Volume Imaging, Hilversum, Netherlands), as described previously (20). TUNEL, PROTEOSTAT, and DAPI positive cells were quantified from 40× confocal microscopy images (317 × 317 μm) using ImageJ (version 1.53n) (41), as described previously (7). Co-labeling of nuclei by TUNEL and PROTESOTAT was quantified from 100× confocal microscopy images (2 regions of 127 × 127 μm) using the Coloc 2 plugin in Fiji (version 2.1.0/1.53c) (42), as described previously (20).

### Characterization of aggregation in vitro in HEK293 cells

DNA constructs coding for murine, human, and bovine rhodopsin with a yellow fluorescent protein (YFP) variant or mTurquoise2 (mTq2), both tagged with a 1D4 epitope, were described previously (7, 32, 43, 44). The K296E mutation was introduced into each of these constructs adapting procedures in the QuickChange II Site-Directed Mutagenesis Kit (Agilent Technologies, Santa Clara, CA) using the following forward and reverse primers: murine rhodopsin, 5’-ACTCTGCCAGCTTTCTTTGCTGAGAGCTCTTCCA and 5’-TGGAAGAGCTCTCAGCAAAGAAAGCTGGCAGAGT; human rhodopsin, 5’-CAGCGTTCTTTGCCGAGAGCGCCGCCATC and 5’-GATGGCGGCGCTCTCGGCAAAGAACGCTG; bovine rhodopsin 5’-CCGGCTTTCTTTGCCGAGACTTCTGCCGTCT and 5’-AGACGGCAGAAGTCTCGGCAAAGAAAGCCGG. HEK293T/17 cells (Cat. No. CRL-11268, American Type Culture Collection, Manassas, VA) were cotransfected with constructs coding for the YFP-and mTq2-tagged rhodopsin and FRET assay conducted on a FluoroMax-4 spectrofluorometer (Horiba Jobin Yvon, Edison, NJ), as described previously (17). Total, *n*-dodecyl-*β*-D-maltoside (DM)-sensitive, and DM-insensitive FRET signals were computed and FRET curves generated by plotting the FRET efficiency versus the acceptor:donor (A:D) ratio and fitting the data by non-linear regression to a rectangular hyperbolic function using Prism 10 (GraphPad Software, San Diego, CA): *E* = (*E*_max_ × A:D)/(EC_50_ + A:D) (17). The non-specific FRET *E*_max_ was defined previously (44).

### Statistics

All statistical analyses were conducted using Prism 10 (GraphPad Software, San Diego, CA), including ANOVA and post-hoc analysis and extra sum of squares F tests.

## Data availability

All data supporting the findings of this study are available within the paper and in supplementary information files. Raw data are available from the corresponding author upon reasonable request.

## Author contributions

S.V. and V.P. conducted experiments and edited the manuscript. S.V., V.P., and P.S.-H.P. designed experiments and analyzed data. P.S.-H.P. wrote the manuscript.

## Acknowledgements

We thank John Denker for generating DNA constructs, genotyping mice, sequencing DNA, testing sgRNA for cutting efficiency and validating knockin mice, Heather Butler for breeding and maintaining mouse colonies, Dawn Smith for culturing HEK293 cells and generating cryosections, Catherine Doller for generating cryosections, and Maryanne Pendergast for help with microscopy and processing of images. We thank the Case Transgenic and Targeting Facility and the Genomics Core at Case Western Reserve University School of Medicine (Cleveland, OH) for generating and identifying K296E rhodopsin knockin mice. This work was funded by grants from the National Institutes of Health (R01EY021731, P30EY011373, and UL1RR024989).

## References

1. Meng, D., Ragi, S. D., and Tsang, S. H. (2020) Therapy in Rhodopsin-Mediated Autosomal Dominant Retinitis Pigmentosa. Mol. Ther. 28, 2139–2149

2. Hartong, D. T., Berson, E. L., and Dryja, T. P. (2006) Retinitis pigmentosa. Lancet 368, 1795–1809

3. Athanasiou, D., Aguila, M., Bellingham, J., Li, W., McCulley, C., Reeves, P. J., and Cheetham, M. E. (2018) The molecular and cellular basis of rhodopsin retinitis pigmentosa reveals potential strategies for therapy. Prog. Retin. Eye Res. 62, 1–23

4. Krebs, M. P., Holden, D. C., Joshi, P., Clark, C. L., 3rd, Lee, A. H., and Kaushal, S. (2010) Molecular mechanisms of rhodopsin retinitis pigmentosa and the efficacy of pharmacological rescue. J. Mol. Biol. 395, 1063–1078

5. Gal, A., ApfelstedtSylla, E., Janecke, A. R., and Zrenner, E. (1997) Rhodopsin mutations in inherited retinal dystrophies and dysfunctions. Prog. Retin. Eye Res. 16, 51–79

6. Dryja, T. P., McGee, T. L., Reichel, E., Hahn, L. B., Cowley, G. S., Yandell, D. W., Sandberg, M. A., and Berson, E. L. (1990) A point mutation of the rhodopsin gene in one form of retinitis pigmentosa. Nature 343, 364–366

7. Vasudevan, S., Senapati, S., Pendergast, M., and Park, P. S. (2024) Aggregation of rhodopsin mutants in mouse models of autosomal dominant retinitis pigmentosa. Nat Commun 15, 1451

8. Park, P. S. (2014) Constitutively active rhodopsin and retinal disease. Adv. Pharmacol. 70, 1–36

9. Keen, T. J., Inglehearn, C. F., Lester, D. H., Bashir, R., Jay, M., Bird, A. C., Jay, B., and Bhattacharya, S. S. (1991) Autosomal dominant retinitis pigmentosa: four new mutations in rhodopsin, one of them in the retinal attachment site. Genomics 11, 199–205

10. Owens, S. L., Fitzke, F. W., Inglehearn, C. F., Jay, M., Keen, T. J., Arden, G. B., Bhattacharya, S. S., and Bird, A. C. (1994) Ocular manifestations in autosomal dominant retinitis pigmentosa with a Lys-296-Glu rhodopsin mutation at the retinal binding site. Br. J. Ophthalmol. 78, 353–358

11. Robinson, P. R., Cohen, G. B., Zhukovsky, E. A., and Oprian, D. D. (1992) Constitutively active mutants of rhodopsin. Neuron 9, 719–725

12. Chen, J., Shi, G., Concepcion, F. A., Xie, G., Oprian, D., and Chen, J. (2006) Stable rhodopsin/arrestin complex leads to retinal degeneration in a transgenic mouse model of autosomal dominant retinitis pigmentosa. J. Neurosci. 26, 11929–11937

13. Li, T., Franson, W. K., Gordon, J. W., Berson, E. L., and Dryja, T. P. (1995) Constitutive activation of phototransduction by K296E opsin is not a cause of photoreceptor degeneration. Proc. Natl. Acad. Sci. U. S. A. 92, 3551–3555

14. Brill, E., Malanson, K. M., Radu, R. A., Boukharov, N. V., Wang, Z., Chung, H. Y., Lloyd, M. B., Bok, D., Travis, G. H., Obin, M., and Lem, J. (2007) A novel form of transducin-dependent retinal degeneration: accelerated retinal degeneration in the absence of rod transducin. Invest. Ophthalmol. Vis. Sci. 48, 5445–5453

15. Chiang, W. C., Hiramatsu, N., Messah, C., Kroeger, H., and Lin, J. H. (2012) Selective activation of ATF6 and PERK endoplasmic reticulum stress signaling pathways prevent mutant rhodopsin accumulation. Invest. Ophthalmol. Vis. Sci. 53, 7159–7166

16. Saliba, R. S., Munro, P. M., Luthert, P. J., and Cheetham, M. E. (2002) The cellular fate of mutant rhodopsin: quality control, degradation and aggresome formation. J. Cell Sci. 115, 2907–2918

17. Gragg, M., and Park, P. S. (2019) Detection of misfolded rhodopsin aggregates in cells by Forster resonance energy transfer. Methods Cell Biol. 149, 87–105

18. Sakami, S., Maeda, T., Bereta, G., Okano, K., Golczak, M., Sumaroka, A., Roman, A. J., Cideciyan, A. V., Jacobson, S. G., and Palczewski, K. (2011) Probing mechanisms of photoreceptor degeneration in a new mouse model of the common form of autosomal dominant retinitis pigmentosa due to P23H opsin mutations. J. Biol. Chem. 286, 10551–10567

19. Hicks, D., and Molday, R. S. (1986) Differential immunogold-dextran labeling of bovine and frog rod and cone cells using monoclonal antibodies against bovine rhodopsin. Exp. Eye Res. 42, 55–71

20. Vasudevan, S., and Park, P. S. (2025) A Y178C rhodopsin mutation causes aggregation and comparatively severe retinal degeneration. Cell Death Discov 11, 32

21. Shen, D., Coleman, J., Chan, E., Nicholson, T. P., Dai, L., Sheppard, P. W., and Patton, W. F. (2011) Novel cell-and tissue-based assays for detecting misfolded and aggregated protein accumulation within aggresomes and inclusion bodies. Cell Biochem. Biophys. 60, 173–185

22. Sieving, P. A., Fowler, M. L., Bush, R. A., Machida, S., Calvert, P. D., Green, D. G., Makino, C. L., and McHenry, C. L. (2001) Constitutive “light” adaptation in rods from G90D rhodopsin: a mechanism for human congenital nightblindness without rod cell loss. J. Neurosci. 21, 5449–5460

23. Sieving, P. A., Richards, J. E., Naarendorp, F., Bingham, E. L., Scott, K., and Alpern, M. (1995) Dark-light: model for nightblindness from the human rhodopsin Gly-90-->Asp mutation. Proc. Natl. Acad. Sci. U. S. A. 92, 880–884

24. Rao, V. R., Cohen, G. B., and Oprian, D. D. (1994) Rhodopsin mutation G90D and a molecular mechanism for congenital night blindness. Nature 367, 639–642

25. Colozo, A. T., Vasudevan, S., and Park, P. S. (2020) Retinal degeneration in mice expressing the constitutively active G90D rhodopsin mutant. Hum. Mol. Genet. 29, 881–891

26. Kobal, N., Krasovec, T., Sustar, M., Volk, M., Peterlin, B., Hawlina, M., and Fakin, A. (2021) Stationary and Progressive Phenotypes Caused by the p.G90D Mutation in Rhodopsin Gene. Int J Mol Sci 22

27. Kremers, G. J., Goedhart, J., van Munster, E. B., and Gadella, T. W., Jr. (2006) Cyan and yellow super fluorescent proteins with improved brightness, protein folding, and FRET Forster radius. Biochemistry 45, 6570–6580

28. Goedhart, J., von Stetten, D., Noirclerc-Savoye, M., Lelimousin, M., Joosen, L., Hink, M. A., van Weeren, L., Gadella, T. W., Jr., and Royant, A. (2012) Structure-guided evolution of cyan fluorescent proteins towards a quantum yield of 93%. Nat Commun 3, 751

29. Park, P. S. (2021) Supramolecular organization of rhodopsin in rod photoreceptor cell membranes. Pflugers Arch. 473, 1361–1376

30. Gragg, M., Kim, T. G., Howell, S., and Park, P. S. (2016) Wild-type opsin does not aggregate with a misfolded opsin mutant. Biochim. Biophys. Acta 1858, 1850–1859

31. Kaushal, S., and Khorana, H. G. (1994) Structure and function in rhodopsin. 7. Point mutations associated with autosomal dominant retinitis pigmentosa. Biochemistry 33, 6121–6128

32. Vasudevan, S., and Park, P. S. (2021) Differential Aggregation Properties of Mutant Human and Bovine Rhodopsin. Biochemistry 60, 6–18

33. Kawamura, S., Colozo, A. T., Muller, D. J., and Park, P. S. (2010) Conservation of molecular interactions stabilizing bovine and mouse rhodopsin. Biochemistry 49, 10412–10420

34. Azam, M., and Jastrzebska, B. (2025) Mechanisms of Rhodopsin-Related Inherited Retinal Degeneration and Pharmacological Treatment Strategies. Cells 14

35. Kari, S., Subramanian, K., Altomonte, I. A., Murugesan, A., Yli-Harja, O., and Kandhavelu, M. (2022) Programmed cell death detection methods: a systematic review and a categorical comparison. Apoptosis 27, 482–508

36. Silverman, D., Chai, Z., Yue, W. W. S., Ramisetty, S. K., Bekshe Lokappa, S., Sakai, K., Frederiksen, R., Bina, P., Tsang, S. H., Yamashita, T., Chen, J., and Yau, K. W. (2020) Dark noise and retinal degeneration from D190N-rhodopsin. Proc. Natl. Acad. Sci. U. S. A. 117, 23033–23043

37. Kwasnieski, J. C., Mogno, I., Myers, C. A., Corbo, J. C., and Cohen, B. A. (2012) Complex effects of nucleotide variants in a mammalian cis-regulatory element. Proc. Natl. Acad. Sci. U. S. A. 109, 19498–19503

38. Senapati, S., Gragg, M., Samuels, I. S., Parmar, V. M., Maeda, A., and Park, P. S. (2018) Effect of dietary docosahexaenoic acid on rhodopsin content and packing in photoreceptor cell membranes. Biochim. Biophys. Acta 1860, 1403–1413

39. Molday, R. S., and MacKenzie, D. (1983) Monoclonal antibodies to rhodopsin: characterization, cross-reactivity, and application as structural probes. Biochemistry 22, 653–660

40. Vasudevan, S., Samuels, I. S., and Park, P. S. (2023) Gpr75 knockout mice display age-dependent cone photoreceptor cell loss. J. Neurochem. 167, 538–555

41. Schneider, C. A., Rasband, W. S., and Eliceiri, K. W. (2012) NIH Image to ImageJ: 25 years of image analysis. Nat. Methods 9, 671–675

42. Schindelin, J., Arganda-Carreras, I., Frise, E., Kaynig, V., Longair, M., Pietzsch, T., Preibisch, S., Rueden, C., Saalfeld, S., Schmid, B., Tinevez, J. Y., White, D. J., Hartenstein, V., Eliceiri, K., Tomancak, P., and Cardona, A. (2012) Fiji: an open-source platform for biological-image analysis. Nat. Methods 9, 676-682

43. Miller, L. M., Gragg, M., Kim, T. G., and Park, P. S. (2015) Misfolded opsin mutants display elevated beta-sheet structure. FEBS Lett. 589, 3119–3125

44. Gragg, M., and Park, P. S. (2018) Misfolded rhodopsin mutants display variable aggregation properties. Biochim. Biophys. Acta 1864, 2938–2948

