## Supplemental Figures and Tables for "Aggregation of the constitutively active K296E rhodopsin mutant contributes to retinal degeneration"

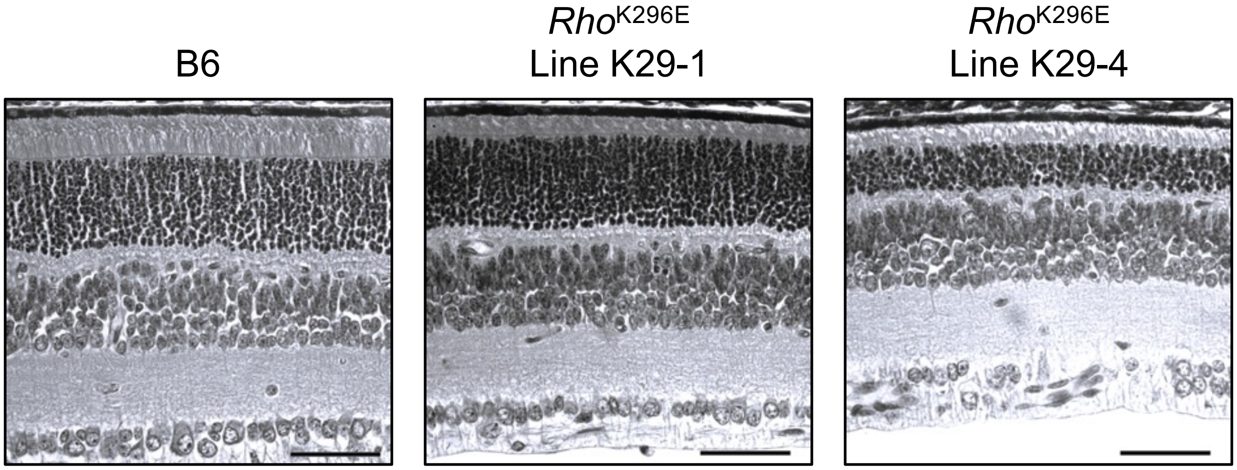


**Figure 1. Different severity of retinal degeneration in *Rho*$^{K296E}$ mice.** Images of retinal sections from 2-week-old B6 mice and *Rho*$^{K296E}$ mice from lines K29-1 and K29-4 are shown. Scale bar, 50 μm.

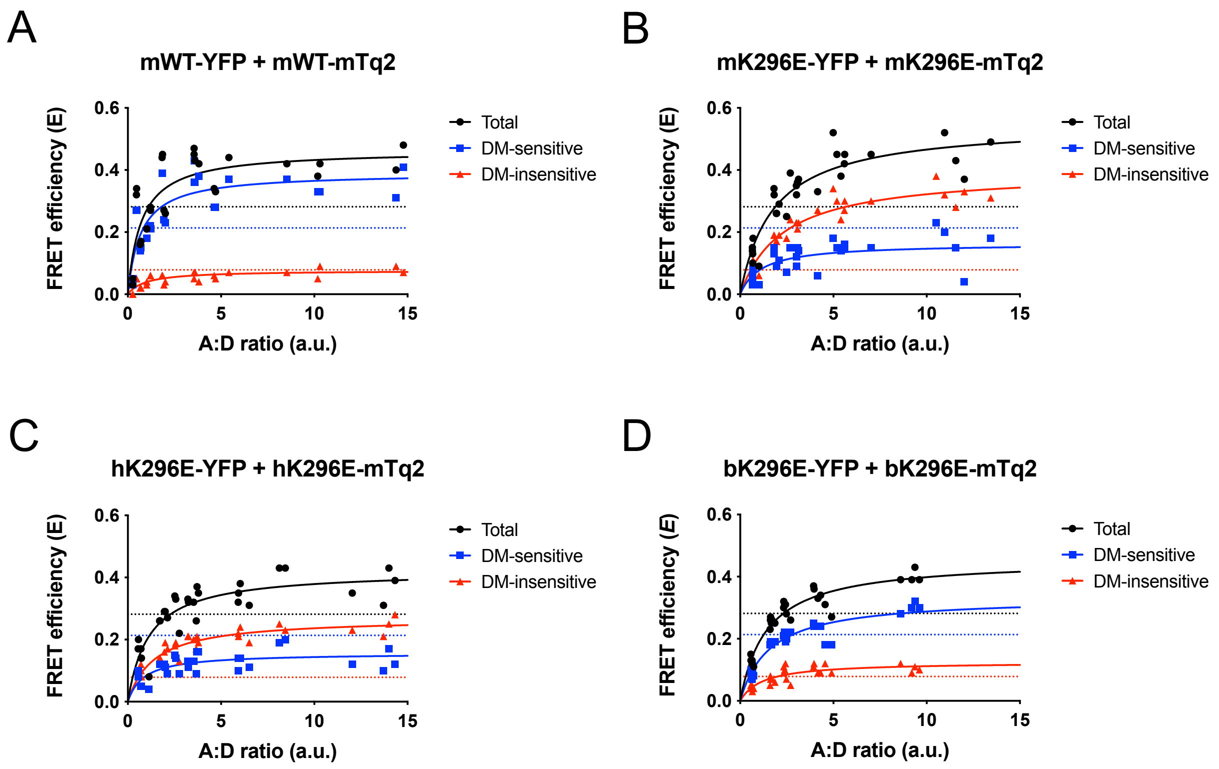


**Figure 2. FRET curves**. FRET curves were generated from cells expressing the indicated YFP-tagged and mTq2-tagged murine WT, murine K296E, human K296E, or bovine K296E rhodopsin. Total (black), DM-sensitive (blue), and DM-insensitive (red) FRET curves are shown. Each curve contains data from 6 separate experiments, which were simultaneously fit with a rectangular hyperbolic function. Fitted lines are shown and values obtained from fits are reported in Figs. 7A and 7B and Supplementary Table 1. The non-specific $E_{\max}$, defined previously (1), is indicated by the dashed lines.

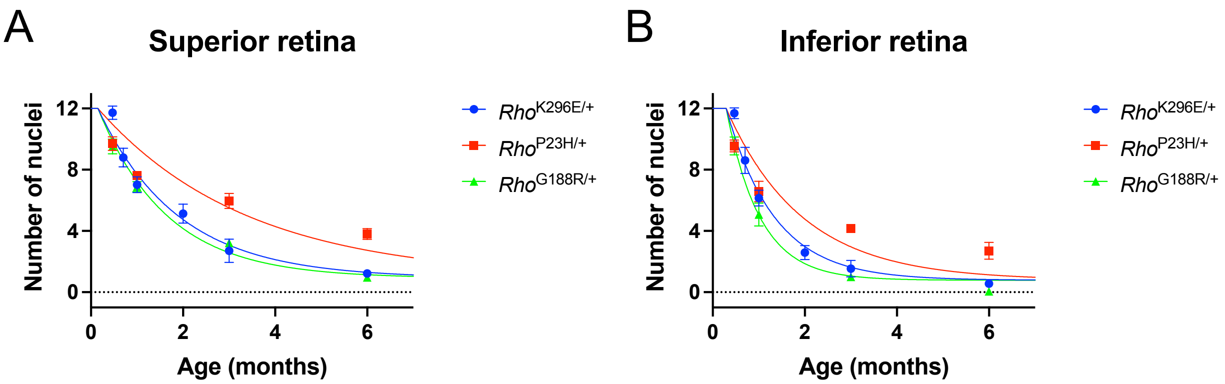


**Figure 3. Kinetics of photoreceptor cell loss in *Rho*$^{K296E/+}$, *Rho*$^{P23H/+}$, and *Rho*$^{G188R/+}$ mice.** Data for *Rho*$^{K296E/+}$ mice (from Fig. 4C) are combined with data for *Rho*$^{P23H/+}$ and *Rho*$^{G188R/+}$ mice (published previously (2)) to visualize the differences in the rate of photoreceptor cell loss among the mouse lines in the superior (A) and inferior (B) retina. Data were fit to a model for plateau followed by one phase exponential decay. The parameter $y_0$ was fixed at 12 and the *plateau* was fit to be common among data sets.

**Table 1.** Kinetics of photoreceptor cell loss. Fitted parameters from Fig. 4C are shown with the standard errors.

| Region of retina | $k$ (month$^{-1}$) | | $x_o$ (month) | |
|---|---|---|---|---|
| | ***Rho***$^{\mathbf{K296E/+}}$ | ***Rho***$^{\mathbf{K296E}}$ | ***Rho***$^{\mathbf{K296E/+}}$ | ***Rho***$^{\mathbf{K296E}}$ |
| **Superior** | 0.66 ± 0.04 | 11.15 ± 3.89 | 0.31 ± 0.04 | 0.38 ± 0.03 |
| **Inferior** | 1.23 ± 0.07 | 7.70 ± 1.79 | 0.43 ± 0.02 | 0.33 ± 0.03 |

Data were fit to a model for plateau followed by one phase exponential decay. The parameter $y_0$ was fixed at 12 and the *plateau* was fit to be common among data sets (0.73 ± 0.11 nuclei). An extra sum of squares F test indicated that the superior and inferior data were significantly different for *Rho*$^{K296E/+}$ mice and required different fitted curves ($P < 0.0001$, $F$ (2, 67) = 24.50) whereas the superior and inferior data were similar for *Rho*$^{K296E}$ mice and could be equally fit by a single curve ($P = 0.3125$, $F$ (2, 43) = 1.195).

**Table 2.** FRET curve analysis

| Coexpressed receptors and treatment | Parameter | | | | | |
|---|---|---|---|---|---|---|
| | $E_{max}$ | | | EC$_{50}$ | | |
| | **Total** | **DM-sensitive** | **DM-insensitive** | **Total** | **DM-sensitive** | **DM-insensitive** |
| **mWT-YFP + mWT-mTq2** | 0.46 ± 0.04 | 0.39 ± 0.03 | 0.08 ± 0.01 | 0.72 ± 0.21 | 0.70 ± 0.22 | 0.95 ± 0.35 |
| **mK296E-YFP + mK296E-mTq2** | 0.55 ± 0.04 | 0.16 ± 0.02 | 0.40 ± 0.02 | 1.80 ± 0.37 | 1.14 ± 0.57 | 2.30 ± 0.31 |
| **hK296E-YFP + hK296E-mTq2** | 0.42 ± 0.02 | 0.16 ± 0.01 | 0.27 ± 0.01 | 1.14 ± 0.24 | 0.86 ± 0.31 | 1.40 ± 0.25 |
| **bK296E-YFP + bK296E-mTq2** | 0.46 ± 0.02 | 0.33 ± 0.02 | 0.13 ± 0.01 | 1.43 ± 0.18 | 1.57 ± 0.24 | 1.19 ± 0.32 |

Data in Supplementary Fig. 2 was fit to a rectangular hyperbolic function as described in the Methods. Fitted values for the maximal FRET efficiency ($E_{max}$) and EC$_{50}$ are shown along with the standard errors.

### References

1. Gragg, M., and Park, P. S. (2018) Misfolded rhodopsin mutants display variable aggregation properties. *Biochim. Biophys. Acta* **1864**, 2938-2948

2. Vasudevan, S., Senapati, S., Pendergast, M., and Park, P. S. (2024) Aggregation of rhodopsin mutants in mouse models of autosomal dominant retinitis pigmentosa. *Nat. Commun.* **15**, 1451
